# Experimental assessment of 3D-printed traps and chemical attractants for the collection of wild *Drosophila melanogaster*

**DOI:** 10.1101/2025.01.28.635319

**Authors:** Alexandra H. Keene-Snickers, Tillie J. Dunham, Mark D. Stenglein

## Abstract

*Drosophila melanogaster*, the common fruit fly, has been instrumental to our understanding of evolution, genetics and disease. There are benefits to studying these flies in the wild, including assessment of their naturally occurring microbiota. To facilitate efforts to catch wild *D. melanogaster*, we designed two fly traps and evaluated several candidate attractants. The first trap utilized a stable food substrate that can be used to catch live flies to establish new lab colonies. The second trap was designed to be reusable and easy to ship to enable the collection of flies over time from diverse locations. We evaluated several chemical attractants derived from banana and from marula fruit, which is the proposed ancestral food host of *D. melanogaster*. We found that wild flies were preferentially attracted to banana-based odorants over marula-derived ones. Overall, these traps and attractants represent an inexpensive and simple option for the collection of wild *D. melanogaster* and related species for sampling or colony establishment.

## INTRODUCTION

Research involving *Drosophila melanogaster* has led to major advances in our understanding of genetics, molecular and cellular biology, and disease^1^. Although laboratory fly lines are useful, neither their genetics nor the laboratory environment necessarily recapitulate the situation in the wild. Studies of *Drosophila* evolution, microbiome, and virus infection have benefited from the use of wild-caught flies^2–5^. The study of the wild fly microbiota reflects an increasing general interest in the natural microbiota of model organisms. For instance, viruses isolated from wild-caught *Caenorhabditis* nematodes have enabled studies of small interfering RNA responses to natural virus infection^6^. Wilding mice, created by implanting laboratory mouse embryos into wild mothers, exhibit more natural immune response^7,8^. Similarly, wild *D. melanogaster* harbor diverse microbes not necessarily present in laboratory strains. These interact with the host and other microbiota, altering host biology and evolution^2,9–12^. The capture and maintenance of wild derived flies in the lab has been instrumental to understanding the impact of common *D. melanogaster* infecting viruses^2,3,13–16^.

To capture wild *D. melanogaster*, researchers typically use manual or electronic aspirators, net swipes, or baited traps consisting of fruit (commonly banana) and yeast^17–20^. Although these traps are effective at capturing wild flies, they involve specialized equipment or could be technically challenging for non-experts^17,20^. Other trap types aim to attract and kill pest *Drosophila* species. *Drosophila suzukii*, a major agricultural pest of soft fruits can be efficiently captured and killed using vinegar-based traps^21–23^. The attractiveness of a trap is important to how successful it will be, particularly in natural environments where there is competition from other food sources.

In this study, we experimentally assessed the stability and attractiveness of different bait substrates, trap designs, and banana- and marula-derived attractants. The marula fruit is proposed to have been an important host fruit for ancestral *D. melanogaster* ^*24*^. Populations of

*D. melanogaster* in Sub-Saharan Africa have polymorphisms in the Or22a-expressing olfactory sensory neurons that increase their sensitivity to ethyl isovalerate, a compound in marula fruit^24^. Other compounds in marula fruit include isoamyl acetate, which is also a main component of synthetic banana flavoring.

Our goal was to design traps that gave us the ability to collect and retain live flies if we wanted and to be easily used by participatory scientists for crowdsourced fly collection. To this end we designed two traps. The first trap focused on the capture of live flies for the generation of new lab colonies. This trap relied on an environmentally stable cornmeal-based food and the addition of isoamyl acetate to increase attractiveness. Our second trap was designed to be easily used by citizen scientists. This trap used banana and yeast, a maximally attractive and readily available bait. These traps could also be reused making them useful for longitudinal collections.

## METHODS AND MATERIALS

### Fly Food

Cornmeal-based food: We used the standard cornmeal food based recipe from the Bloomington Drosophila Stock Center (https://bdsc.indiana.edu/information/recipes/bloomfood.html). Twenty-five to 50 mL of food was placed in fly bottes (VWR, 75813-140). For experiments involving ethyl isovalerate, ethyl isovalerate (Sigma-Aldrich 112283) was added to final percent volume concentrations of 0.45%, 0.9%, 1.8%, 3.6% or 7.2%. For experiments involving isoamyl acetate, synthetic banana flavoring (Kroger Brand Imitation Banana, 87-9110) was added to a final concentration of 1.5%.

Potato-based food: was prepared as described by Waddle, 1996 (www.anapsid.org/fruitfly.html), except baker’s yeast was substituted for brewer’s yeast^25^. Briefly, 120g of instant potato was mixed with 12g yeast. Mold inhibitor was made by adding 1g of Tegosept to 1 L of boiling water. Mold inhibitor was added to the potato/yeast mixture to rehydrate the potatoes. The volume of mold inhibitor varied from batch to batch based on how much liquid was needed to fully rehydrate the potato/yeast mixture. Twenty-five to 50 mL of food was then placed in fly bottles and sprinkled with baker’s yeast. For attractiveness experiments, 4g of agar dissolved in 200mL water was added to increase shelf life.

Banana-based food: was prepared as described by Hundt, 1996 (www.anapsid.org/fruitfly.html)^25^. Briefly, 4 bananas were mashed together with 1/8 cup cane sugar until liquified. Rolled oats were added until the mixture became firm. Twenty-five to 50 mL of food was transferred to fly bottles and sprinkled with baker’s yeast.

### Fly food attractants

All potential attractants except the homemade banana extract were obtained through commercial vendors: isoamyl acetate (Kroger Brand Imitation Banana, 87-9110), ethyl isovalerate (Sigma-Aldrich, 112283) and marula oil (Rocky Mountain Oil Company, All Organic marula Oil). Homemade banana extract was made by manually mashing bananas until mostly liquid. The pulp and liquid was then separated using a mesh strainer. The liquid was retained and stored in a -20°C freezer until use.

### Outdoor experiments

For attractiveness experiments, wild-derived flies from our lab’s FoCo17 colony were used^26^. Microcosms were constructed using bug dorms (dimensions: W32.5 x D32.5 x H77.0 cm; similar to: bugdorm, BD4F3074). Microcosms were placed in an outdoor environment that experienced mixed sun and shade with temperatures ranging from 13°C to 38°C. Microcosms were protected from rain by a tarp. Flies released into microcosms were given 24-48 hours to select one of the foods, either plain or supplemented with one of the marula fruit or banana additives. To trap the flies, we used food bottles capped with 3D printed trap lids. These lids allowed flies to easily enter the bottle but not as easily get out. Bottles were collected and frozen before the number of flies in each bottle were counted.

### Indoor experiments

Population cages in our insectary were used for these experiments. These cages were approximately D1.8 x W1.2 x H2.4 m in size. Procedures were the same as for the outdoor experiments: food was placed in bottles with the trap lid, flies were released for 24-48 hours before food bottles were collected, frozen and the number of flies in each bottle was determined. Indoor microcosms were on a 12-hour light-dark cycle and temperature was kept at ∼21°C.

### Live collection trap

3D-printed trap components were designed in the OpenSCAD scripting design language and printed in polylactic acid on a LulzBot Taz6 printer (Fargo Additive Manufacturing Equipment). Designs are available at https://github.com/stenglein-lab/3D_parts/tree/master/fly_trap. For our initial trap testing, we used threadless fly bottles (VWR, 75813-140) and attached the trap lid to the bottle using parafilm. Subsequent experiments involving volunteer collectors used 4 oz jars (Uline, S-9934) and the 3D printed trap containing cornmeal-based food as described above.

Before pouring the food into bottles, we added isoamyl acetate at a 1:100 ratio to food. Fifty mL of food was poured into bottles and left to cool overnight. Bottles were stored at 4°C until use. Each bottle was sealed with a cotton plug and parafilm. Traps were shipped to volunteers along with the 3D printed trap lid and parafilm. Volunteers were instructed to place the trap lid on the bottle and place the trap in a location with fruit fly activity (one outdoor location and one indoor location). After ∼1 week volunteers were instructed to seal the bottle with the provided parafilm and to ship it back to our lab (**Fig. 4A**).

### Reusable trap

We created kits containing materials for trapping and returning flies. Kits included a clear plastic jar (Uline, S-9934), a 3D printed trap lid, a 3D printed barrier with organdy mesh (Oriole Textile, 2060), Fleischmann’s Instant Dry Yeast, a screw top 2.0 mL tube and a pre-paid shipping return envelope. Volunteers were instructed to place 1-2 slices of banana in the bottom of the jar with a small pinch of yeast. The barrier was placed over the banana/yeast ensuring that flies remained separated from bait. Volunteers were instructed to replace the banana/yeast every 3-5 days depending on the environmental conditions (hot/humid environments were told to replace the banana/yeast more often) (**Fig. 4B**). Flies were collected from traps that had been placed in a - 20°C freezer for ≥ 30 minutes. Once dead, flies could be removed by tapping the jar upside down or with forceps and stored frozen at -20°C in 2.0 mL tubes. Tubes were returned in the provided prepaid shipping envelope.

### Data analyses

Data visualization and statistical analyses were performed in RStudio (v 2024.09.0+375) using the tidyverse package^27^. Analysis code is available at https://github.com/LKeene/fly_food. A p-value of α = 0.05 or lower was used as our significance threshold. For attractiveness experiment, negative binomial models were generated using the MASS package and comparisons were made using the emmeans package^28,29^. Pairwise comparisons were adjusted for multiple hypothesis testing using Tukey’s method. For Figure 3C, a pseudocount of one was added to all values to accommodate zero counts.

## RESULTS

### Cornmeal-based fly food was most stable

We first set out to assess the stability of traps baited with different food substrates over two weeks. We tested cornmeal-, potato-, and banana-based foods, all of which are commonly used to rear *D. melanogaster*. Each type was made plain or with synthetic banana flavoring (isoamyl acetate). Food bottles were placed in a controlled indoor environment or outside for two weeks. Bottles were visually assessed daily to identify signs of decay and microbial growth.

Banana-based food was the least stable with visible mold and bacterial growth occurring in a matter of days (**Supplemental Fig. 1**). Potato-based food was more stable: after two weeks there was visible mold and bacterial growth but less than the banana-based food (**Supplemental Fig. 1**). Cornmeal-based food was the most stable in indoor and outdoor environments (**Supplemental Fig 1**). The indoor cornmeal-based food looked nearly the same at the end of the two weeks as it did at the start. The presence of synthetic banana flavoring did not impact the stability of any of the foods. For all food types there was considerable evaporation in the bottles that caused moisture to accumulate, particularly outdoors (**Supplemental Fig. 1**).

**Supplemental Figure 1:**
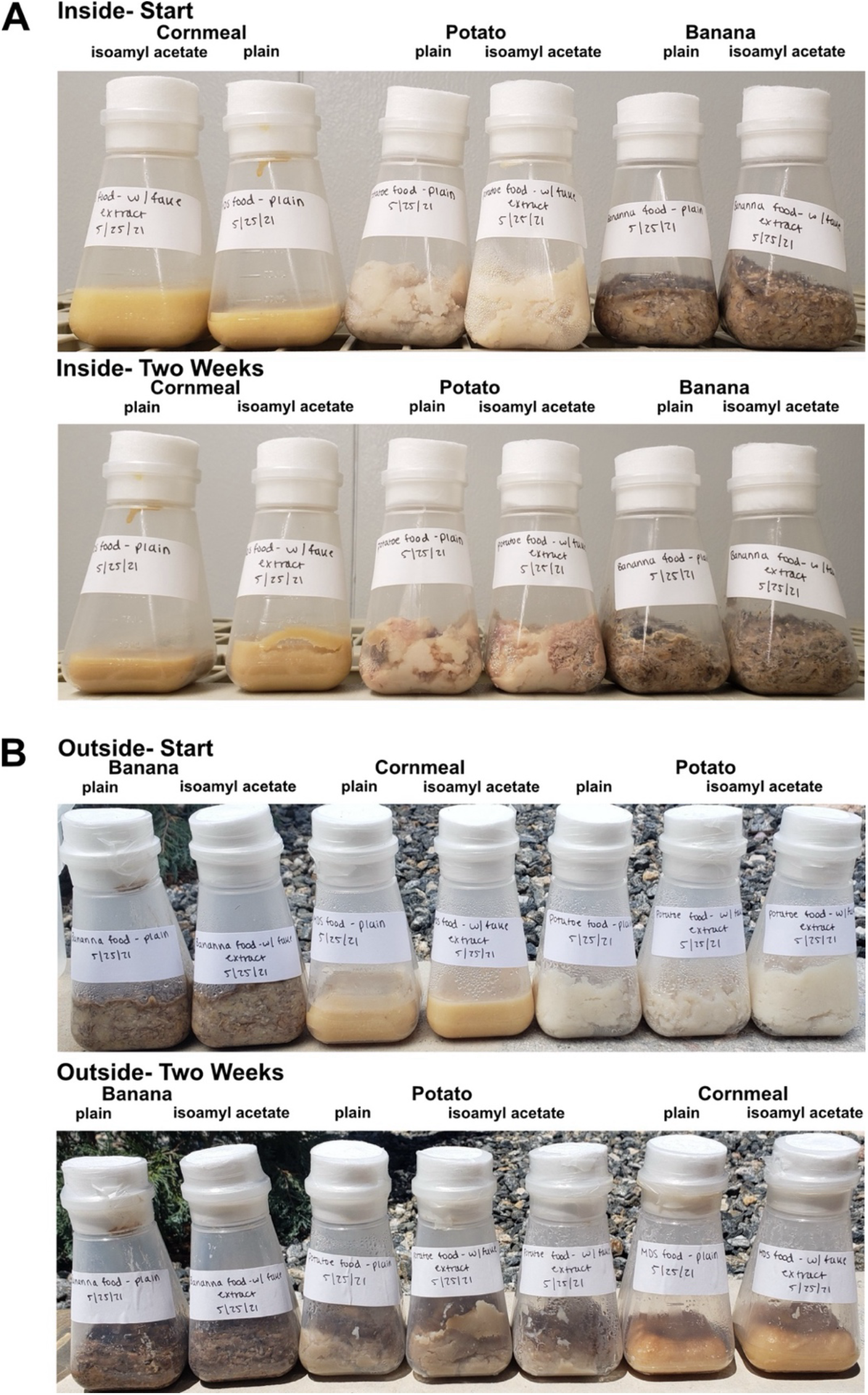
Cornmeal-based fly food was stable over two weeks in indoor and outdoor environments. Cornmeal-, potato-, and banana-based fly foods were placed either in our indoor insectary (A) or outside (B). Representative before and after pictures are shown for both experiments. The food was visually inspected daily over the experimental period.

### Banana-based compounds increase the attractiveness of plain food to *D. melanogaster*

We next sought to augment the attractiveness of cornmeal food baited traps. Although flies can be maintained on such food in the lab, it was unclear whether these foods would be sufficiently attractive. We expected that there would be competition from other food sources, particularly in outdoor environments, where organic volatiles known to attract *D. melanogaster* are emitted from rotting plant debris^30,31^. *D. melanogaster* are attracted to fermenting banana catalyzed by the addition of yeast^32^. However, this banana and yeast mixture decays over the course of several days creating a sticky substance that makes it difficult to retrieve flies. We therefore employed a choice assay with our wild derived laboratory line of *D. melanogaster* (FoCo-17) where the flies were exposed to plain cornmeal-based food with or without homemade banana extract, or to banana/yeast bait^26^.

We released flies into mesh enclosures in controlled indoor or outdoor environments containing traps baited with different food types. We left flies for 24-48 hours and recorded the number of flies in each bottle. Flies preferred food with a banana component (**Fig. 1**). Banana and yeast food was more attractive than either plain cornmeal (p = 1.7×10^-10^) or cornmeal food supplemented with homemade banana extract (p = 5.4×10^-3^). This preference was less pronounced in outdoor conditions (p = 8.8×10^-3^ & p = 0.83, **Fig. 1**). This shows that banana and yeast was an effective fly attractant, as expected. However, a banana-based additive also increased attractiveness.

**Figure 1.**
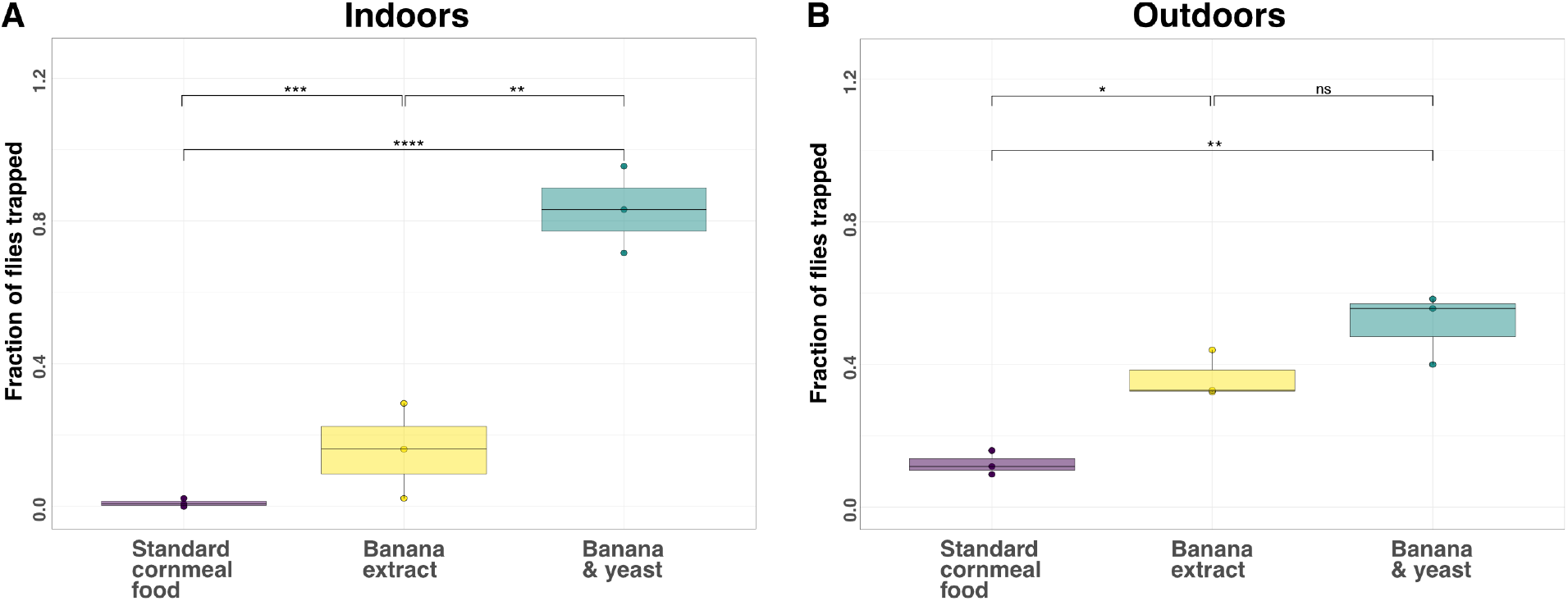
*D. melanogaster* were more attracted to banana-based baits. Fly bottles were either filled with plain cornmeal-based food, cornmeal-based food with homemade banana extract or banana and yeast. A 3D printed fly trap lid was used to seal the bottles. FoCo-17 flies were released in controlled environments either inside (n = 143, n = 197, n = 173) or outside (n=173, n=61, n=220) and left for 24-48 hours. Bottles were collected and the proportion of flies in each bottle was tabulated. Significance values from a negative binomial model are: p < 0.0001 ‘****’, 0.0001 < p > 0.001 ‘***’, 0.001 < p > 0.01 ‘**’, 0.01 < p > 0.05 ‘*’ and p > 0.05 ‘ns’.

### Banana extract was more attractive to *D. melanogaster* than marula tree derivatives

The fruit of the marula tree has been proposed to be the ancestral host of *D. melanogaster*^24^. We therefore tested whether ethyl isovalerate, a volatile ester emitted from marula fruit, would be better at attracting flies than banana-based compounds. Southern African *D. melanogaster* populations have polymorphisms in the Or22a odorant receptor that increase sensitivity to this compound, but this has not yet been tested using North American populations^24^. Since flies were highly attracted to banana, we also tested the use of synthetic banana flavoring (isoamyl acetate) as an inexpensive and readily available chemical attractant. Isoamyl acetate is also a prominent ester in marula fruit^24,33^.

We also wanted to test whether the effectiveness of these additives was impacted by the food base. Fly bottles with cornmeal or potato food and the various attractants were placed in indoor or outdoor enclosures. Flies from the wild-derived FoCo-17 population were released for 24-48 hours and the number of flies in each trap was recorded.

Across all replicates in both environments regardless of food type, banana-based odorants were the most attractive (**Fig. 2**). Surprisingly, there were very few flies in the ethyl isovalerate or marula fruit oil bottles (**Fig. 2**). There was a decrease in the relative attractiveness of banana extracts outdoors (**Fig. 2B & D**). Standard cornmeal-based food with either homemade or store-bought banana extract produced statistically indistinguishable results (**Fig. 2**). Due to the ease of use and chemical consistency, we opted to use synthetic banana flavoring (isoamyl acetate) for subsequent experiments.

**Figure 2:**
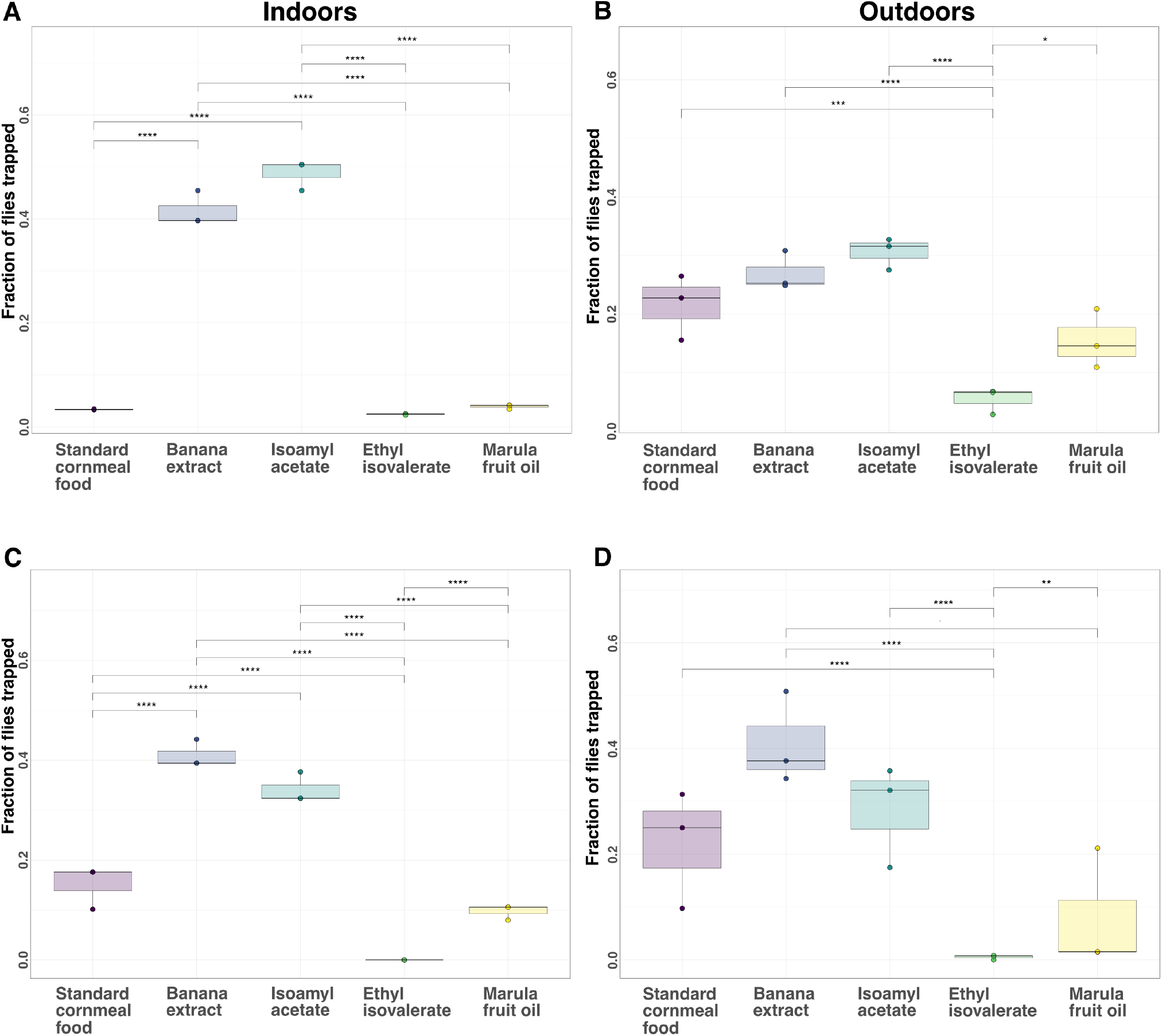
*D. melanogaster* prefer food with a banana-based attractant over food with derivatives of the marula fruit. The relative attractiveness of cornmeal-based (A & B) or potato-based (C & D) fly food with various baits were assessed in competition experiments. Experiments were conducted indoors (A,C) and outdoors (B,D). FoCo-17 flies were released for 24-48 hours in enclosures containing baited fly traps. The number of flies in each replicate was: (A) indoor n= 121, 121, 88; (B) outdoor n = 211, 206, 378 and (C) indoor n = 142, 142, 138; (D) outdoor n = 134, 246, 260. Significance values determined from a negative binomial model: p < 0.0001 ‘****’, 0.0001 < p > 0.001 ‘***’, 0.001 < p > 0.01 ‘**’, 0.01 < p > 0.05 ‘*’ and p > 0.05 ‘ns’.

The lack of attractiveness of ethyl isovalerate led us to test whether we were using an appropriate concentration. We performed an experiment using five ethyl isovalerate concentrations: 0.45%, 0.9%, 1.8% (∼the natural concentration in marula fruit^33^), 3.6% and 7.2%. Plain cornmeal bottles and bottles with cornmeal plus isoamyl acetate at 1.5%, the natural concentration found in marula fruit^33^, served as controls. As with the previous experiments, FoCo-17 flies were released in our indoor mesocosm for 24-48 hours and then counted to determine the proportion of flies in each bottle.

Flies were most attracted to plain food or food containing synthetic banana extract (**Fig. 3**). The ethyl isovalerate bottles in aggregate attracted on average 31.8% of the flies with the 1.8% ethyl isovalerate bottle attracting 10.5% of the flies. The 3.6% and 7.2% bottles attracted the fewest flies. In contrast, synthetic banana flavoring attracted on average 37.4% of the flies and plain food 30.6% of the flies (**Fig. 3**). Food with banana extract was not significantly more attractive than plain food. It is possible that the plain and banana extract food bottles were not placed far enough apart due to the limited space in the mesocosm. Overall, the results of these experiments confirmed our previous findings that *D. melanogaster* were more attracted to isoamyl acetate than ethyl isovalerate.

**Figure 3:**
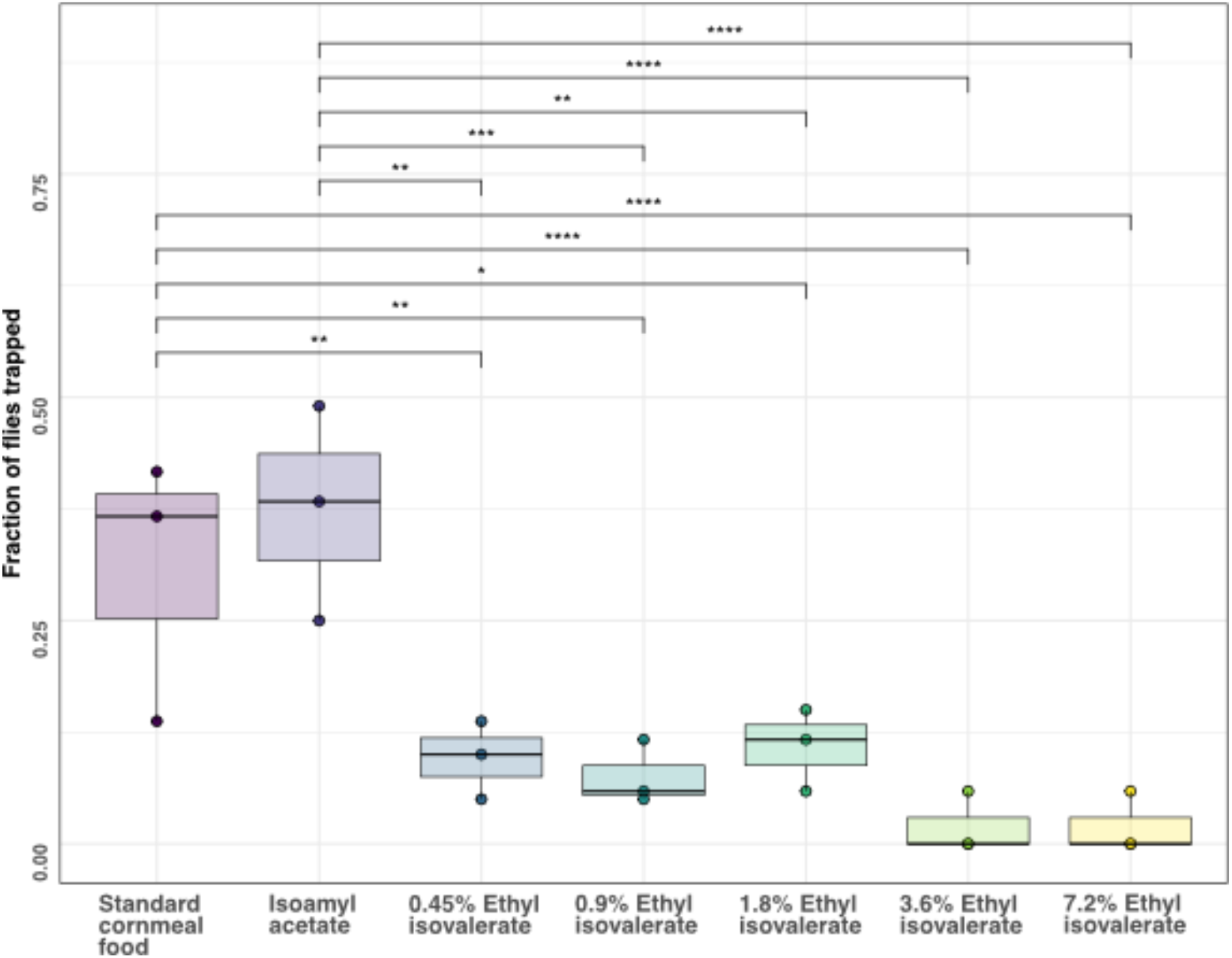
An outbred population of *D. melanogaster* from Colorado was not attracted to ethyl isovalerate. Flies from the FoCo-17 wild-derived outbred population of *D. melanogaster* were released (n=51, 60, 60) in an indoor microcosm cage for 24-48 hours. Cornmeal-based food with the following additives were placed in each cage: plain food, 1.5% synthetic banana extract and food supplemented with 0.45%, 0.9%, 1.8%, 3.6% or 7.2% ethyl isovalerate. After 24-48 hours the number of flies were counted. Significance values determined from pairwise comparisons of a negative binomial model are: p < 0.0001 ‘****’, 0.0001 < p > 0.001 ‘***’, 0.001 < p > 0.01 ‘**’, 0.01 < p > 0.05 ‘*’ and p > 0.05 ‘ns’.

### A kit for capture of wild *D. melanogaster*

As a pilot test of the ability of these traps to be used by non-specialists, we sent traps with cornmeal-based food with isoamyl acetate to three volunteers across the United States (**Fig. 4A**). Volunteers were instructed to place the traps in an outdoor or indoor location with fruit fly activity for roughly a week. They were told to watch for trapped flies and signs of larval development. Once first instar larva could be seen, the volunteers were instructed to seal the bottle with parafilm and ship it back to our lab. We received bottles from two locations and were able to generate colonies that persist three years later.

**Figure 4:**
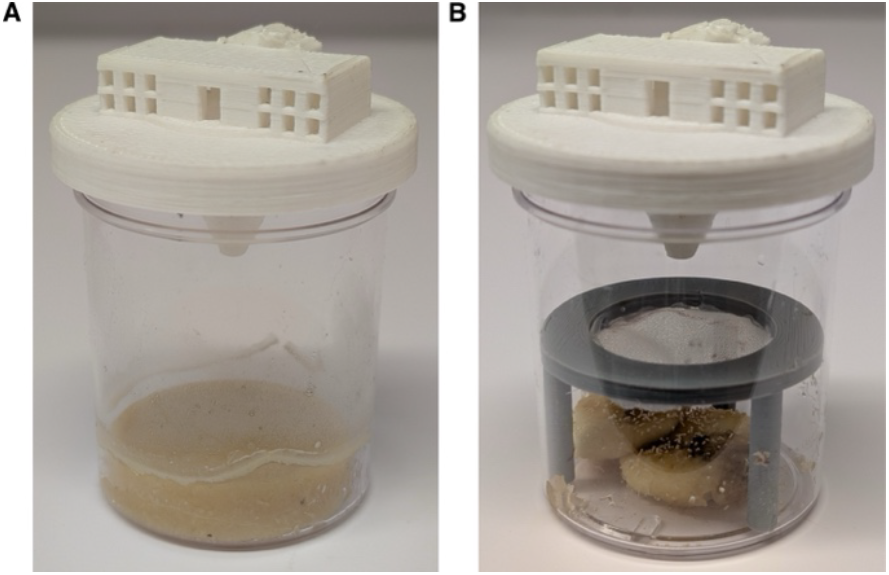
A 3D printed fly trap kit to collect wild *D. melanogaster*. Fly trap designs for the collection of live flies (A) or for repeated fly collection over time (B). Trap components are 3D printed. Cornmeal-based food with isoamyl acetate can be used as a stable food that facilitates egg laying (A) or banana and yeast can be used as bait without the risk of flies becoming trapped in the decomposing fruit (B).

We next tested a modified trap that simplified fly capture and trap reuse. This trap included a 3D printed “table” covered in fine mesh to separate flies from the bait (**Fig. 4B**). This trap design takes advantage of the attractiveness of banana/yeast bait while preventing flies from getting entrapped.

These traps proved to be an effective means to capture flies. Volunteers were instructed to add one to two slices of banana and a small pinch of yeast to the bottom of the jar, place the “table” over the banana and close the jar with the trap lid. They then placed the trap in an area with fruit fly activity and monitored the trap for flies and rotting of the bait. Volunteers periodically froze the trap to kill the flies and moved them to the 2.0 mL tube. The tube was stored in the freezer and another round of collection began with fresh bait. At the end of the collection period (1-3 months) they shipped the 2.0 mL tubes back to our lab. Once at our lab, fly species was determined using morphological characteristics and stored in a -80°C freezer until further analysis.

## DISCUSSION

Here we describe the design and effectiveness of two wild *D. melanogaster* fly traps. The first trap is characterized by a stable cornmeal and agar-based food supplemented with propionic acid to inhibit bacterial growth. Eggs laid by adult flies remained viable in this food even after shipment. This made it possible to generate new laboratory colonies from various locations after collection by non-specialists. We have maintained two stocks generated from flies collected in this manner in 2021.

The second trap was designed to facilitate sample collection and trap reuse. This makes it ideal for crowdsourced sample collection and longitudinal collection. One feature of this trap is that it can be combined with other baits for the collection of specific *Drosophila* species^20^. In a small field trial, kits were sent to nine volunteers with no previous *Drosophila* collecting experience. Volunteers were able to setup the kits and three locations successfully collected flies that were received by our laboratory.

We tested the attractiveness of marula fruit oil and ethyl isovalerate as a *D. melanogaster* attractants because it had been proposed that ancestral populations of these flies evolved to specialize on marula fruit^24^. However, we found that banana-based additives were more attractive, at least to our wild-derived flies from Colorado (**Fig. 2 & Fig. 3**). Although *D. melanogaster* are considered to be generalists, many species within the *melanogaster* subgroup specialize on a single fruit with populations of the same species showing different specializations^24,34,35^. It is possible that the population of *D. melanogaster* we tested have either lost sensitivity to ethyl isovalerate or gained sensitivity to volatiles released by different rotting fruits^24,31^. This could explain the observed unattractiveness of this compound. There is considerable variation in odorant receptors that detect esters emitted by yeast and fermenting fruit ^36^.

The use of community engagement to further scientific research has become increasingly popular^37–39^. The traps are designed for use by untrained volunteers and to be baited with readily available materials. *D. melanogaster* are attracted to a variety of other fruits if banana is unavailable^40^. Overall, our flexible designs enable researchers and non-specialists to obtain flies from diverse locations for minimal cost and effort.

## ACKNOWLEDGEMENTS

We thank volunteers and their willingness to participate in trap testing: Shaun Cross, Brendan, Arabella, and Malila Kavaney, Amanda, Mike, Clara, and Zoe Furlotti, Lisa Harrington, Paul Keene, Khwannarin Khemsom, and Anastazia Jablunovsky. We would also like to thank Keller Gonzales for technical assistance.

## REFERENCES

1. Yamaguchi, M. & Yoshida, H. Drosophila as a Model Organism. in Advances in Experimental Medicine and Biology vol. 1076 1–10 (Springer New York LLC, 2018).

2. Martinez, J. et al. Virus evolution in wolbachia-infected drosophila. Proceedings of the Royal Society B: Biological Sciences 286, (2019).

3. Ortiz-Baez, A. S., Shi, M., Hoffmann, A. A. & Holmes, E. C. RNA virome diversity and Wolbachia infection in individual Drosophila simulans flies. J Gen Virol 102, 001639 (2021).

4. Webster, C. L. et al. The Discovery, Distribution, and Evolution of Viruses Associated with Drosophila melanogaster. PLoS Biol 13, e1002210 (2015).

5. Webster, C. L., Longdon, B., Lewis, S. H. & Obbard, D. J. Twenty-Five New Viruses Associated with the Drosophilidae (Diptera). Evolutionary Bioinformatics 12s2, EBO.S39454 (2016).

6. Félix, M.-A. & Wang, D. Natural Viruses of Caenorhabditis Nematodes. Annu Rev Genet 53, 313–326 (2019).

7. Ma, J. et al. Laboratory mice with a wild microbiota generate strong allergic immune responses. Sci Immunol 8, (2023).

8. Rosshart, S. P. et al. Laboratory mice born to wild mice have natural microbiota and model human immune responses. Science (1979) 365, (2019).

9. Shi, M. et al. No detectable effect of Wolbachia wMel on the prevalence and abundance of the RNA virome of Drosophila melanogaster. Proceedings of the Royal Society B: Biological Sciences 285, (2018).

10. Palmer, W. H., Varghese, F. S. & Van Rij, R. P. Natural variation in resistance to virus infection in dipteran insects. Viruses vol. 10 Preprint at 10.3390/v10030118 (2018).

11. Van Rij, R. P. et al. The RNA silencing endonuclease Argonaute 2 mediates specific antiviral immunity in Drosophila melanogaster. Genes Dev 20, 2985–2995 (2006).

12. Nayak, A. et al. Cricket paralysis virus antagonizes Argonaute 2 to modulate antiviral defense in Drosophila. Nat Struct Mol Biol 17, 547–554 (2010).

13. Cogni, R., Ding, S. D., Pimentel, A. C., Day, J. P. & Jiggins, F. M. Wolbachia reduces virus infection in a natural population of Drosophila. Commun Biol 4, 1327 (2021).

14. Wallace, M. A. & Obbard, D. J. Naturally occurring viruses of Drosophila reduce offspring number and lifespan. Proceedings of the Royal Society B: Biological Sciences 291, (2024).

15. Shi, M. et al. No detectable effect of Wolbachia wMel on the prevalence and abundance of the RNA virome of Drosophila melanogaster. Proceedings of the Royal Society B: Biological Sciences 285, (2018).

16. Mackay, T. F. C. et al. The Drosophila melanogaster Genetic Reference Panel. Nature 482, 173–8 (2012).

17. Markow, T. A. & O’Grady, P. M. Collecting Drosophila in the wild. in Drosophila 145–153 (Elsevier, 2006). doi:10.1016/B978-012473052-6/50004-4.

18. Topal, P., Garg, D. & S. Fartyal, R. Fruit Flies ( Drosophila spp. ) Collection, Handling, and Maintenance: Field to Laboratory. in The Wonders of Diptera - Characteristics, Diversity, and Significance for the World’s Ecosystems (IntechOpen, 2021). doi:10.5772/intechopen.97014.

19. Faria, V. G. & Sucena, É. From Nature to the Lab: Establishing Drosophila Resources for Evolutionary Genetics. Front Ecol Evol 5, (2017).

20. Carson, H. L. & Heed, W. B. Methods of Collecting Drosophila. in The Genetics and Biology of Drosophila (eds. Ashburner, M., Thompson, J. N. & Carson H.L.) vol. 3d 1–28 (Academic Press, London, 1983).

21. Landolt, P. J., Adams, T. & Rogg, H. Trapping spotted wing drosophila, Drosophila suzukii (Matsumura) (Diptera: Drosophilidae), with combinations of vinegar and wine, and acetic acid and ethanol. Journal of Applied Entomology 136, 148–154 (2012).

22. Iglesias, L. E., Nyoike, T. W. & Liburd, O. E. Effect of trap design, bait type, and age on captures of Drosophila suzukii (Diptera: Drosophilidae) in berry crops. J Econ Entomol 107, 1508–1512 (2014).

23. Dadour, I. R. & Cook, D. F. THE EFFECTIVENESS OF FOUR COMMERCIAL FLY TRAPS AT CATCHING INSECTS. Aust J Entomol 31, 205–208 (1992).

24. Mansourian, S. et al. Wild African Drosophila melanogaster Are Seasonal Specialists on Marula Fruit. Current Biology 28, 3960-3968.e3 (2018).

25. Kaplan, M., Hundt, A., Waddle, F. & Nehring, N. Homemade fruit Fly Culture media. Melissa Kaplan’s Herp Care Collection (1996).

26. Cross, S. T. et al. Partitiviruses Infecting Drosophila melanogaster and Aedes aegypti Exhibit Efficient Biparental Vertical Transmission. J Virol 94, (2020).

27. Wickham, H. et al. Welcome to the Tidyverse. J Open Source Softw 4, 1686 (2019).

28. Venables, W. & Ripley, B. Modern Applied Statistics with S. (Springer, New York, 2002).

29. Lenth, R. V. et al. emmeans: Estimated Marginal Means, aka Least-Squares Means. CRAN: Contributed Packages Preprint at 10.32614/CRAN.package.emmeans (2017).

30. Ruebenbauer, A., Schlyter, F., Hansson, B. S., Löfstedt, C. & Larsson, M. C. Genetic Variability and Robustness of Host Odor Preference in Drosophila melanogaster. Current Biology 18, 1438–1443 (2008).

31. Soto-Yéber, L., Soto-Ortiz, J., Godoy, P. & Godoy-Herrera, R. The behavior of adult Drosophila in the wild. PLoS One 13, e0209917 (2018).

32. Markow, T. A. & O’Grady, P. M. Chapter 4 - Collecting Drosophila in the wild. in Drosophila (eds. Markow, T. A. & O’Grady, P. M.) 145–153 (Academic Press, San Diego, 2006). doi:10.1016/B978-012473052-6/50004-4.

33. Viljoen, A. M., Kamatou, G. P. P. & Başer, K. H. C. Head-space volatiles of marula (Sclerocarya birrea subsp. caffra). South African Journal of Botany 74, 325–326 (2008).

34. Auer, T. O. et al. Olfactory receptor and circuit evolution promote host specialization. Nature 579, 402–408 (2020).

35. Comeault, A. A. et al. A nonrandom subset of olfactory genes is associated with host preference in the fruit fly Drosophila orena. Evolution Letters vol. 1 73–85 Preprint at 10.1002/evl3.7 (2017).

36. de Bruyne, M., Smart, R., Zammit, E. & Warr, C. G. Functional and molecular evolution of olfactory neurons and receptors for aliphatic esters across the Drosophila genus. J Comp Physiol A Neuroethol Sens Neural Behav Physiol 196, 97–109 (2010).

37. Lukyanenko, R., Wiggins, A. & Rosser, H. K. Citizen Science: An Information Quality Research Frontier. Information Systems Frontiers 22, 961–983 (2020).

38. Fraisl, D. et al. Citizen science in environmental and ecological sciences. Nature Reviews Methods Primers 2, 64 (2022).

39. Encarnação, J., Teodósio, M. A. & Morais, P. Citizen Science and Biological Invasions: A Review. Front Environ Sci 8, (2021).

40. Dweck, H. K. M. et al. The Olfactory Logic behind Fruit Odor Preferences in Larval and Adult Drosophila. Cell Rep 23, 2524–2531 (2018).

